# Proteins associated with *Cryptococcus* spp. extracellular polysaccharides are implicated in virulence, capsule integrity, and biofilm formation

**DOI:** 10.1101/2024.05.14.594069

**Authors:** Piotr R. Stempinski, Ella Jacobs, Livia Liporagi Lopes, Samuel Rodrigues Dos Santos Junior, Kevin Rojas, Maggie P. Wear, Arturo Casadevall

## Abstract

*Cryptococcus neoformans* and *Cryptococcus gattii* are pathogenic fungi comprising species complexes that are the etiological agents of cryptococcosis. Both fungal species complexes possess a variety of virulence factors that include a polysaccharide capsule, melanin production, enzymes, and the release of polysaccharides. In this study, we characterized the proteins associated with cryptococcal polysaccharides and analyzed their function by studying mutants deficient in their expression. Mass spectrometry analysis of exopolysaccharide fractions revealed 62 proteins, many of which were previously uncharacterized. Analysis of capsule size, structure, and integrity in protein-deficient mutants allowed us to identify four proteins, which we named <u>Ex</u>tracellular Polysaccharide <u>R</u>elated proteins (Exa1-4), and whose absence affected proper capsule assembly and EPS release. In summary, cryptococcal exopolysaccharide is associated with proteins that are implicated in virulence, capsular integrity, and biofilm formation. Our results associated new phenotypes for four previously unknown genes and show that the filtered polysaccharide preparations are a rich source of proteins that can inform capsular assembly, virulence, and immunological studies.

## Introduction

The basidiomycetous encapsulated fungi, *Cryptococcus neoformans,* and *Cryptococcus gattii* comprise species complexes that are the two main etiologic agents of life-threatening cryptococcosis [1,2]. The released polysaccharides of *Cryptococcus neoformans* are important virulence factors that play a role in the pathogenicity of this fungus. In particular, glucuronoxylomannan (GXM) and, galactoxylomannan (GalXM), which are the major components of the cryptococcal capsule, are critical for the virulence of *C. neoformans* [3–5]. The cryptococcal capsule is a thick, protective layer that surrounds the cells of the fungus, helping it evade the host immune system and establish infections [6–8]. The capsule plays several important roles in the pathogenesis of *C. neoformans* and *C. gattii* [9–11]. It helps the fungus to evade host immune responses by concealing the epitopes from immune cells and preventing the recognition of proteins located on the cell surface [12–15]. It also protects the fungus from environmental stresses, such as oxidative stress [8]. In addition, the capsule plays a vital role in the process of cryptococcal dissemination in the host tissues, such as the central nervous system [13,16].

The released extracellular polysaccharides (EPS) are also believed to contribute to virulence by interfering with the development of effective immune responses[9]. EPS can interfere with macrophage recognition and phagocytosis, which are important components of the host immune response, allowing the fungus to evade detection and destruction by the immune system [17]. Furthermore, EPS in tissue has been associated with an inadequate inflammatory response, which impairs the ability of the host immune system to clear the fungal infection [17–19]. A filtered EPS preparation known as CneF was extensively studied by Murphy and collaborators in the 1990s and shown to elicit strong T cell response although the antigens responsible for its immunological effects were never characterized, and these were presumably proteins and/or mannoproteins, but none were identified [20,21]. Further characterization of the components found in this fraction could result in the development of potential therapeutics. *C. neoformans* EPS is also an essential component for the formation of biofilm, whereby complex communities of microorganisms can adhere to surfaces and form a protective matrix that helps them resist environmental stresses and host immune responses [22,23]. While biofilms have been implicated in the virulence of many bacterial pathogens, the role of biofilms in the pathogenesis of *Cryptococcus neoformans* infections is less well understood [24,25]. The presence of cryptococcal biofilms on prosthetic joints or medical devices used to treat complications of cryptococcal meningitis such as ventriculoarterial shunts is uncommon but has been described in clinical case reports [26–29]. A better understanding of the biofilm formation requires more extensive research on the function of released proteins and EPS in this process [30,31].

Most studies of capsular structure, architecture, and composition involve the analysis of EPS isolated from the supernatant of cell cultures [9,32]. However, the topic of EPS-related proteins and their possible role in capsule synthesis has not been investigated. In this study, we performed proteomic analysis of EPS from five cryptococcal strains. The selection of two *C. gattii* complex and three *C. neoformans* complex strains allowed us to identify proteins and a potential target for anti-cryptococcal therapeutics. This analysis revealed several proteins, some of which have been associated with the capsule, but many that are associated with other cellular roles. To categorize the proteins, we analyzed the peptide sequence for predicted polysaccharide interacting protein domains. Several of those proteins including Cda1, Cda3, Gox1, Gox2, and Agn1 were previously characterized in different components of *C. neoformans* cells [33,34]. Among the uncharacterized proteins, we identified three with lectin-like domains and one with Glycosyl hydrolase domain GH79, which suggest the role in capsule assembly or EPS release [35,36]. Further investigation allowed us to elucidate the role of previously uncharacterized EPS-related proteins in capsule and biofilm formation and determine their impact on cryptococcal virulence in the *Galleria mellonella* model system [37,38]. Overall, our data suggest that EPS-interacting proteins have important roles important for capsular architecture and biofilm formation which in consequence can dramatically impact cryptococcal virulence. The lack of homology to human proteins suggests that these have promising potential for utilization as drug targets or new antigens for anti-cryptococcal vaccines.

## RESULTS

### Identification of EPS-related proteins

To better understand the composition of proteins associated with EPS we profiled the EPS-associated proteins of three *C. neoformans* complex strains and two *C. gattii* complex strains after placing the cells in capsule-inducing conditions and collecting supernatant of cell cultures [2,39]. Next, the supernatant was fractionated using centrifugation filters (Fig 1). Passage of this material through a 100k Da filter allowed for separation of larger components of supernatant and then a subsequent passage through a 10 kDa filter allowed us to collect and concentrate a fraction of supernatant containing released proteins and EPS. Isolation and concentration of cryptococcal EPS allowed us to visualize the different patterns of released proteins among tested samples by SDS-PAGE (Fig 2). The composition of the released proteins gel pattern was unique for each tested strain. Next, we analyzed the composition of EPS-related proteomes of *C. neoformans* and *C. gattii*. Using mass spectrometry, we detected multiple unique peptides that were then analyzed, and assigned to the corresponding gene ID (Supplementary Table 1). Based on the protein accession number and The Basic Local Alignment Search Tool (BLASTp) analysis we assigned corresponding gene IDs of the four reference cryptococcal strains including *C. neoformans* var*. grubii* H99, *C. neoformans* var. *neoformans* JEC21, *C. gattii* VGII R265 and *C. gattii* VGIII CA1280 [40,41]. For simplification of further analysis, we used gene ID assigned to *C. neoformans* var*. grubii* H99 strain to describe all homologs identified in this study. Among 62 proteins identified based on the unique peptide count, 23 proteins were detected in the samples from both cryptococcal species (Table 1A). Another group of 14 proteins was detected only in EPS obtained from strains that belong to *C. neoformans* complex and 26 proteins only in the *C. gattii* complex (Table 2 and Table 3).

**Fig 1.**
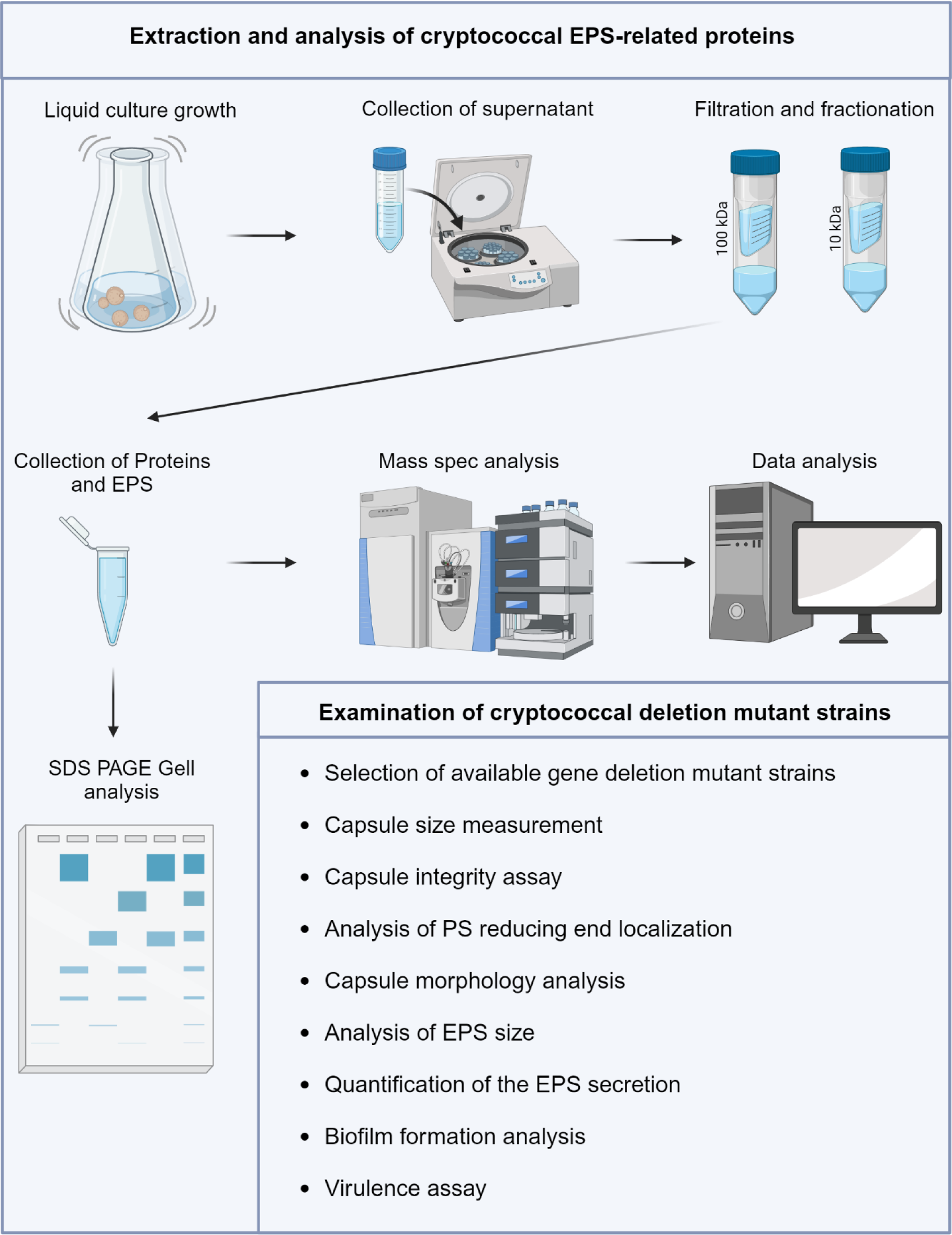
Diagram illustration the process of extraction and analysis of the cryptococcal EPS and EPS -related proteins.

**Fig 2.**
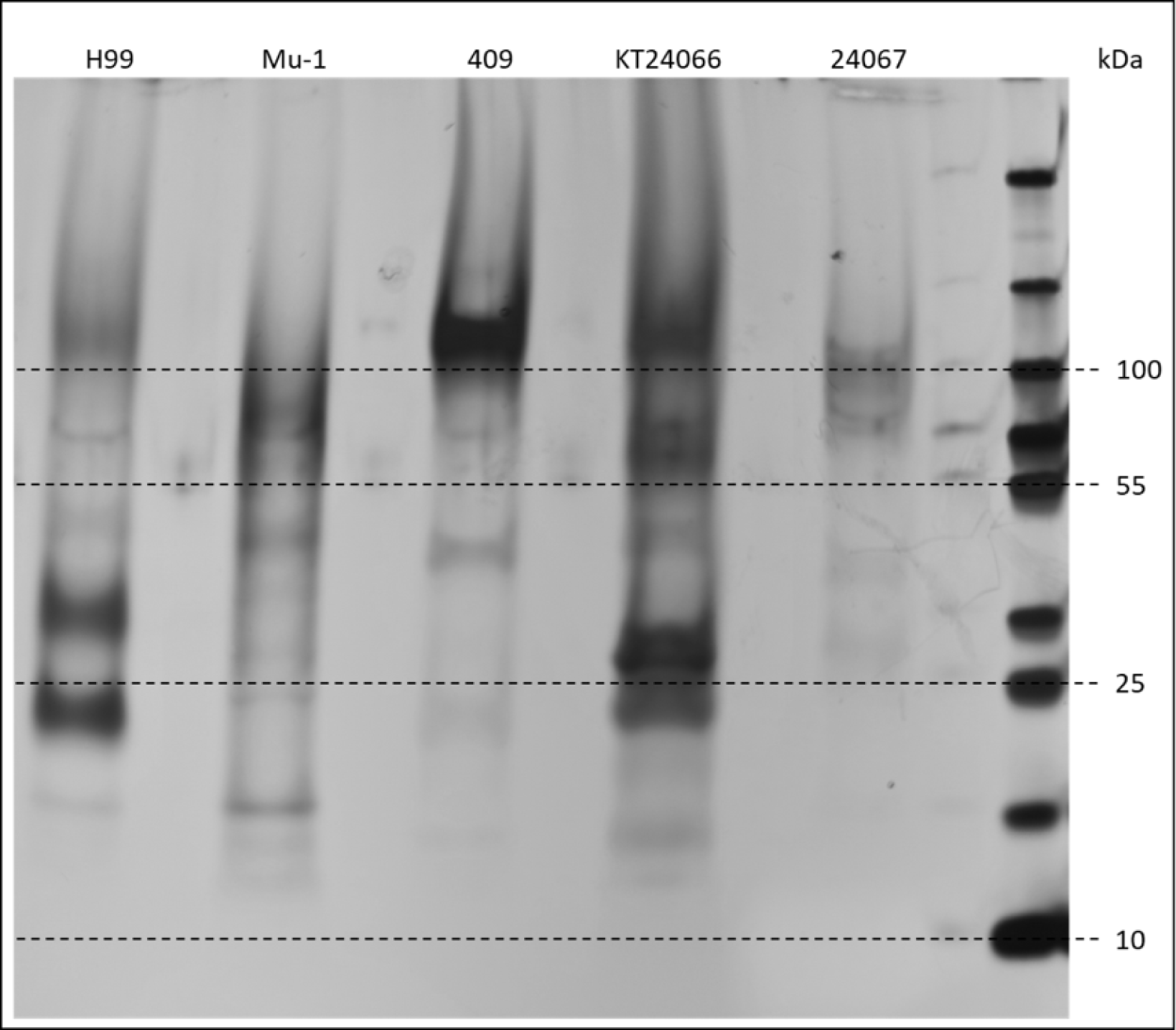
SDS-PAGE analysis of EPS-related proteins among five different cryptococcal strains. Fractionated cryptococcal EPS, containing EPS-related proteins was loaded on 4-20% SDS-PAGE gel. Protein electrophoresis was performed for 2 hours at 60V after which gel was stained with silver stain. Right lane presents protein ladder PageRuler Plus (Thermo scientific).

**Table 1.** Proteins identified in both *C. neoformans* and *C. gattii* EPS.

| *C. neoformans*<br>H99 ID | *C. gattii*<br>R265 ID | Accession | Molecular<br>Mass | Name | Protein |
| --- | --- | --- | --- | --- | --- |
| CNAG_00407 | CNBG_1499 | AFR92540 | 71 kDa | Gox1 | glyoxal oxidase |
| CNAG_00417 | CNBG_1489 | AFR92550 | 47 kDa |  | elongation factor 1-gamma |
| CNAG_00776 | CNBG_1155 | KIR68985 | 38 kDa | Mp88 | immunoreactive mannoprotein MP88 |
| CNAG_00799 | CNBG_1135 | AFR92930 | 85 kDa |  | cellulase |
| CNAG_01019 | CNBG_0599 | AAK31914 | 16 kDa | Sod1 | copper zinc superoxide dismutase |
| CNAG_01047 | CNBG_0624 | KIR49987 | 14 kDa | Exa1 | hypothetical protein |
| CNAG_01239 | CNBG_0806 | AFR94907 | 44 kDa | Cda3 | chitin deacetylase |
| CNAG_01498 | CNBG_3934 | AAW42619 | 68 kDa | Akp2 | Arylsulfatase |
| CNAG_01896 | CNBG_3047 | AFR98092 | 37 kDa |  | alcohol dehydrogenase |
| CNAG_02030 | CNBG_5182 | AFR95790 | 72 kDa | Gox2 | glyoxal oxidase |
| CNAG_02189 | CNBG_5332 | AFR95631 | 57 kDa | Amy1 | alpha-amylase |
| CNAG_02225 | CNBG_5365 | AFR95595 | 49 kDa | Exg104 | glucan 1,3-beta-glucosidase |
| CNAG_02860 | CNBG_3511 | AFR93831 | 49 kDa | Ebg1 | endo-1,3(4)-beta-glucanase |
| CNAG_04869 | CNBG_5798 | AFR97347 | 69 kDa | Pnb1 | carboxylesterase |
| CNAG_04891 | CNBG_5817 | AFR97325 | 22 kDa |  | hypothetical protein |
| CNAG_05312 | CNBG_5735 | AFR94574 | 42 kDa | Vep7 | hypothetical protein |
| CNAG_05799 | CNBG_1745 | AFR96118 | 49 kDa | Cda1 | chitin deacetylase |
| CNAG_05893 | CNBG_1654 | AFR96213 | 38 kDa | Exa2 | hypothetical protein |
| CNAG_06081 | CNBG_4796 | KIR45959 | 92 kDa | Vep13 | glucose oxidase |
| CNAG_06150 | CNBG_4857 | AAN76525 | 78 kDa |  | heat-shock protein 90, partial |
| CNAG_07638 | CNBG_3989 | KGB77902 | 38 kDa | Exa4 | hypothetical protein |
| CNAG_07736 | CNBG_2140 | AFR96695 | 85 kDa | Agn1 | glucan endo-1,3-alpha-glucosidase |

**Table 2.** Proteins identified only in *C. neoformans* EPS.

| *C. neoformans*<br>H99 ID | *C. gattii*<br>R265 ID | Accession | Molecular<br>Mass | Name | Protein |
| --- | --- | --- | --- | --- | --- |
| CNAG_00065 | CNBG_0489 | AFR92203 | 26 kDa |  | ran-binding protein 1 |
| CNAG_01120 | CNBG_9057 | AFR95026 | 51 kDa |  | pyruvate dehydrogenase complex |
| CNAG_01577 | CNBG_3860 | AAW42099 | 49 kDa |  | glutamate dehydrogenase |
| CNAG_01653 | CNBG_3792 | AFR97858 | 30 kDa | Cig1 | cytokine inducing-glycoprotein |
| CNAG_02399 | CNBG_4104 | AFR95434 | 52 kDa | Glr1 | glutathione-disulfide reductase |
| CNAG_02500 | CNBG_4198 | AFR95332 | 61 kDa |  | calnexin |
| CNAG_03007 | CNBG_3648 | AFR93685 | 12 kDa |  | hypothetical protein |
| CNAG_05131 | CNBG_4560 | AFR94396 | 25 kDa |  | hypothetical protein |
| CNAG_05264 | CNBG_5792 | AFR94522 | 64 kDa |  | alpha-amylase AmyA |
| CNAG_05449 | CNBG_0265 | AAW44998 | 13 kDa | Cmt1 | expressed protein |
| CNAG_05918 | CNBG_1632 | ODN80972 | 42 kDa |  | ATP synthase subunit beta, mitochondrial |
| CNAG_06104 | CNBG_4814 | AFR98329 | 71 kDa | Exa3 | hypothetical protein |
| CNAG_06208 | CNBG_4912 | AFR98435 | 86 kDa |  | heat shock 70kDa protein 4 |

**Table 3.** Proteins identified only in *C. gattii* EPS.

| *C. neoformans*<br>H99 ID | *C. gattii*<br>R265 ID | Accession | Molecular<br>Mass | Name | Protein |
| --- | --- | --- | --- | --- | --- |
| CNAG_00256 | CNBG_0315 | ADV20382 | 39 kDa |  | Aspartate-semialdehyde dehydrogenase |
| CNAG_00306 | CNBG_0265 | KIR50366 | 18 kDa | Mtn2 | hypothetical protein |
| CNAG_00587 | CNBG_1361 | KIR69195 | 17 kDa | Ril3 | hypothetical protein |
| CNAG_00588 | CNBG_1362 | ADV19710 | 18 kDa | Ril2 | Hypothetical protein |
| CNAG_00626 | CNBG_1302 | KIR86582 | 26 kDa |  | hypothetical protein |
| CNAG_00699 | CNBG_9112 | KIR69060 | 150 kDa | Lpi15 | transmembrane receptor |
| CNAG_01854 | CNBG_3374 | AFR98049 | 91 kDa | Hep2 | heparinase II/III family protein |
| CNAG_01920 | CNBG_3023 | AAA82978 | 43 kDa | Ubl4 | polyubiquitin |
| CNAG_01993 | CNBG_2959 | KIR37484 | 21 kDa |  | hypothetical protein |
| CNAG_02852 | CNBG_9366 | AFR93840 | 52 kDa |  | 4-aminobutyrate transaminase |
| CNAG_02864 | CNBG_9367 | ALO60484 | 21 kDa |  | hypothetical protein |
| CNAG_02944 | CNBG_3587 | KIR48733 | 59 kDa | Aph1 | vacuolar acid phosphatase |
| CNAG_03413 | CNBG_2195 | ADV23512 | 45 kDa |  | alginate lyase |
| CNAG_03465 | CNBG_2144 | BAG50355 | 67 kDa | Lac1 | diphenol oxidase (Laccase) |
| CNAG_03524 | CNBG_2090 | KIR67909 | 181 kDa |  | transmembrane receptor |
| CNAG_03916 | CNBG_5551 | KIR46581 | 62 kDa |  | glucose-6-phosphate isomerase |
| CNAG_03983 | CNBG_5487 | AFR93483 | 36 kDa |  | oxidoreductase |
| CNAG_04245 | CNBG_2800 | AFR96977 | 86 kDa | Chi22 | chitinase |
| CNAG_04291 | CNBG_2759 | AFR97022 | 36 kDa |  | glycosyl-hydrolase |
| CNAG_05471 | CNBG_4391 | KIR44230 | 108 kDa |  | alpha-glucosidase |
| CNAG_05750 | CNBG_1793 | AAW44019 | 58 kDa |  | ATP synthase alpha chain |
| CNAG_06000 | CNBG_4742 | KIR44370 | 41 kDa |  | glycoprotein |
| CNAG_06125 | CNBG_4834 | AHA60268 | 36 kDa | Tef1 | translation elongation factor EF1-alpha |
| CNAG_06291 | CNBG_5149 | KIR44433 | 27 kDa | Fpd1 | deacetylase |
| CNAG_06501 | CNBG_4970 | AFR98731 | 59 kDa | Gas1 | 1,3-beta-glucanosyltransferase |
| CNAG_06946 | CNBG_0827 | KIR49777 | 38 kDa |  | hypothetical protein |
| CNAG_07004 | CNBG_5871 | AAN75618 | 54 kDa | Lpd1 | dihydrolipoyl dehydrogenase |

Previous observations of cryptococcal Extracellular Vesicles (EVs) indicated that polysaccharides, including GXM and proteins, can be exported via EVs [42–45]. To analyze if EVs-dependent transportation plays a major role in the secretion of proteins found in EPS isolates we confronted our results with previous EVs proteomic projects [43–45]. A comparison of the proteins identified in our study with those reported in three proteomic studies of EVs-related proteins indicated that 35 (56.5%) of the proteins from our set were previously detected in cryptococcal EVs (Supplementary Table 2).

To examine the potential function of the detected proteins with polysaccharide metabolism, we analyzed their sequences for catalytic and carbohydrate-binding functional domains using the Carbohydrate Active Enzymes database (CAZy)[46]. Among the 62 proteins detected in our study, 22 had structural motifs with possible carbohydrate enzymatic activity (Fig 3A). The most common carbohydrate-acting protein domain was Carbohydrate-Binding Module Family 13 (CBM13). CBM13 was identified in the sequence of a putative cellulase and three proteins with unknown function were detected in our study. This domain has been previously found in multiple proteins, including plant lectins, with the ability to bind mannose and xylan, which correlates with the main components of cryptococcal GXM [47–50]. Another carbohydrate interacting domain commonly found in our study is Glycoside Hydrolase Family 5 (GH5), previously known as cellulase family A. This family includes multiple proteins with endoglucanase, exoglucanases, endomannanases, or exomannanases activity [50,51].

**Fig 3.**
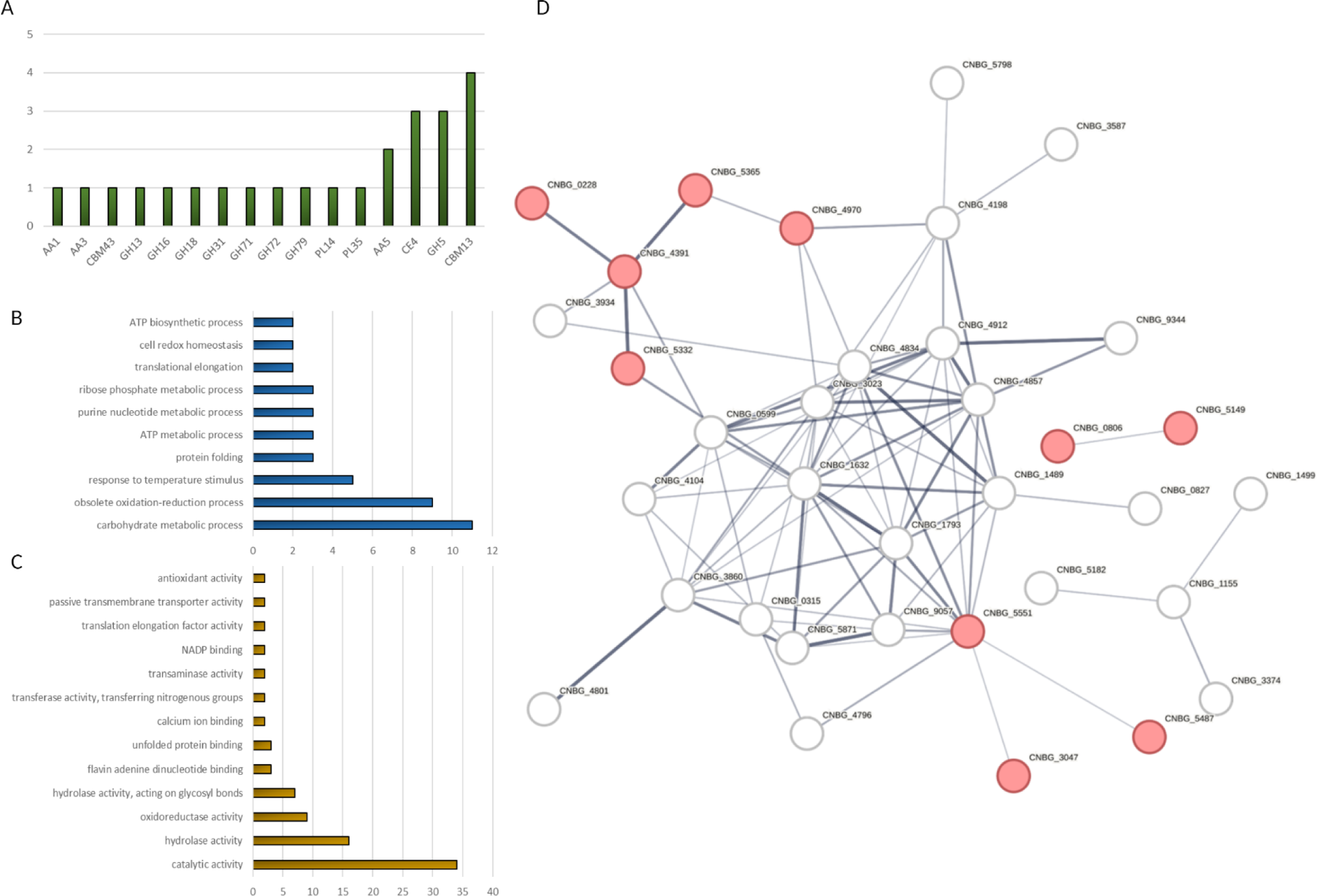
(**A**) Number of carbohydrates interacting domains detected among all EPS-related proteins found in this study. The y-axis refers to the number of proteins found in cryptococcal EPS with the respective protein domains listed on the X-axis. (**B**) GO enrichment analysis of the proteins identified in the EPS collected from *C. neoformans* and *C. gattii*. in biological processes. (**C**) GO enrichment analysis of the proteins identified in the EPS collected from *C. neoformans* and *C. gattii* in molecular function of the protein. The y-axis refers to GO terms, the x-axis describes the numbers of proteins identified in the EPS collected matching the GO category. (**D**) Graphical presentation of protein interaction network among the EPS-related cryptococcal proteins. The network was organized and arranged using STRING protein network (VER 11.5) software. Proteins without confirmed protein-protein interaction are not displayed in the network. Red nodules indicate established proteins involved in carbohydrate metabolism process.

Gene ontology (GO) analysis showed that the most represented proteins in the group were involved in carbohydrate metabolic processes (Fig 3B). Proteins involved in the oxidation-reduction process and response to temperature stimuli were also common in the analyzed proteins. GO analysis for molecular function revealed that the most significantly represented processes involved catalytic activity, hydrolase activity, and oxidoreductase activity, respectively (Fig 3C). To visualize the interaction between proteins detected in cryptococcal EPS samples we created a graphical presentation of protein-protein interaction network using STRING software (Fig 3D). Many of the detected proteins were previously reported to interact with other EPS-related proteins directly or indirectly.

Protein sequence analysis with the BLASTP tool was used to determine the homology of cryptococcal EPS-related proteins to human proteins and proteins of major fungal pathogens (*Candida albicans*, *Histoplasma capsulatum,* and *Aspergillus fumigatus*)[52–56]. The majority of the analyzed proteins presented very low (<40%) or no homology to proteins found in human cells which indicates them as promising novel anti-fungal drug targets or vaccine antigens[57–59]. Among the analyzed proteins 16 presented no significant homology to proteins found in other major fungal pathogens indicating their potential cryptococcus-specific function (Supplementary Table 3).

The cell wall and polysaccharide capsule are the two most peripheral structures of the cryptococcal cells [60–62]. The cell wall plays an essential role in the organization, maintenance, and support of the structure of the polysaccharide capsule [60,62]. The computational analysis of the predicted GPI-anchor sites in the released EPS-associated cryptococcal proteins detected in our study revealed that 16 had a predicted GPI-anchor in their structure. Among these, 14 had a high GPI-anchor prediction score suggesting a connection between capsular and extracellular released proteins with the structure of the cell wall (Supplementary Table 3).

### Role of the uncharacterized EPS-related proteins in capsule maintenance and assembly

To identify novel proteins involved in the creation and maintenance of cryptococcal capsule or regulation of EPS secretion we focused on those with uncharacterized functions (Table 2). For further analysis, we selected 11 proteins that were currently represented in the single gene deletion collection. These strains were analyzed for cell body size and capsule diameter after placement in capsule-inducing media and compared to their parental strain. To better evaluate the influence of single gene deletion on capsule size we calculated the ratio of capsule diameter to cell diameter (Fig 4). Based on our observation strains deficient in three genes (CNAG_01047, CNAG_05893, CNAG_06104) each were associated with a significant decrease in capsule size. In contrast, deficiency in CNAG_07638 was associated with a significant increase in capsule size. Three genes for which deletion caused a decrease in capsule size (CNAG_01047, CNAG_05893, CNAG_06104) were named <u>Ex</u>tracellular Polysaccharide <u>R</u>elated proteins *exa1*, *exa2*, and *exa3* respectively. The gene associated with the mutant exhibiting enlarged capsule size (CNAG_07638) was named *exa4*.

**Fig 4.**
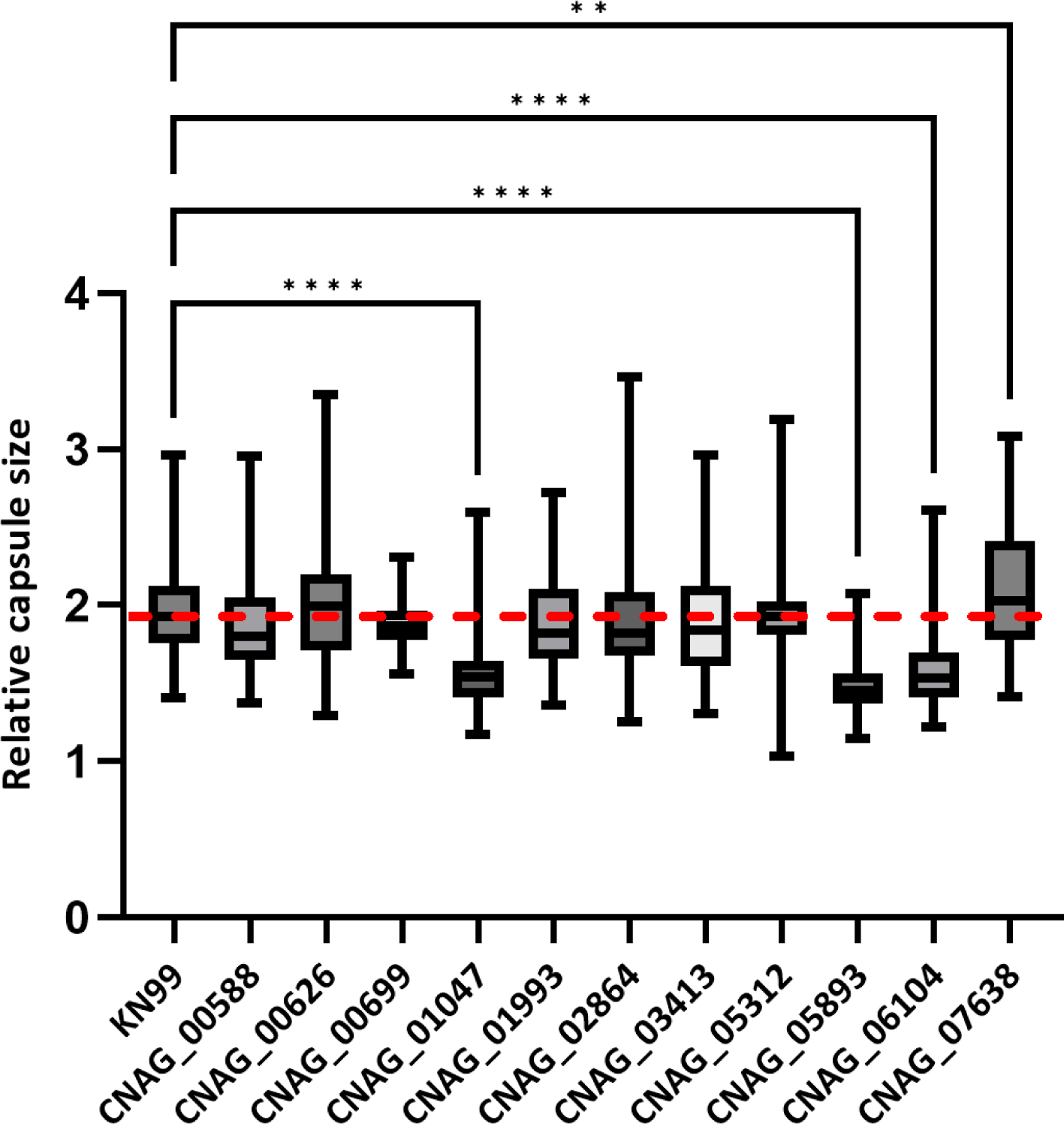
Capsular characterization of selected single gene mutant strains. (A) Capsule size analysis of India ink-stained cryptococcal cells quantified by automated quantitative capsule analysis (QCA) by comparing ratio of capsule diameter to cell diameter. The red dotted line indicates the average ratio value for WT strain. (B) Visualization of cells with alternated capsule size. Capsule size alternation was analyzed by ANOVA on set of >180 cells, (* p < 0.05, ** p < 0.05, ∗∗∗p < 0.001). The red dotted line indicates average, relative capsule size for WT strain.

The capsule structure of selected cryptococcal deletion mutant strains that exhibited enlarged or decreased capsule size was further analyzed using a capsule integrity assay based on the reduction in capsule size in cells after exposure to physical stress created with a horn-sonicator (Fig 5). Two of the deletion mutant strains manifested a significantly changed capsule size compared to the stressed WT cells. Despite the initially enlarged capsule, *exa4*Δ cells exhibited the most drastic reduction of capsule integrity (73.9% of the original ratio) resulting in a capsule size significantly smaller than in the WT cells. On the contrary, the *exa2*Δ strain was the only strain with an initially reduced capsule size that presented a significantly smaller capsule size when compared to the WT strain after the sonication stress test. Sonicated *exa2*Δ cells presented the smallest capsule size among all tested strains suggesting the essential role of this protein in the maintenance of capsule integrity.

**Fig 5.**
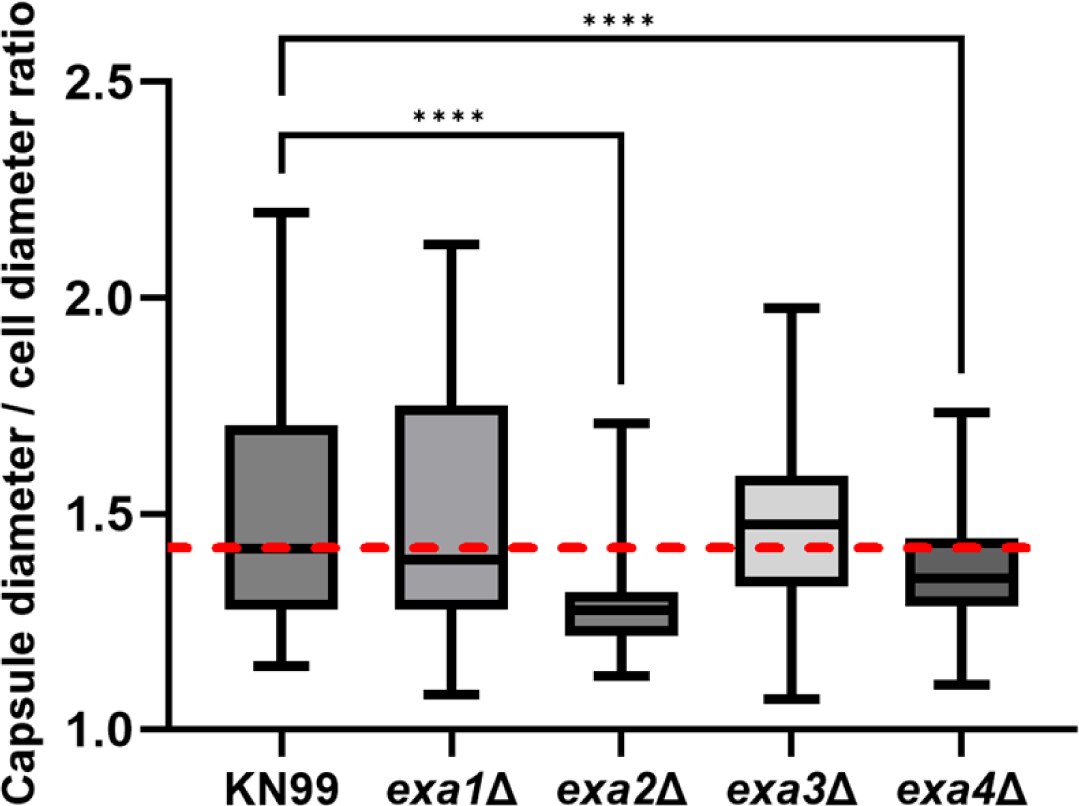
Capsule integrity assay of selected single gene mutant strains. Capsular integrity was measured in encapsulated cells exposed to 15 sec of sonication with 17W power output. Capsule size analysis of India ink-stained cryptococcal cells quantified by automated quantitative capsule analysis (QCA) by comparing the ratio of capsule diameter to cell diameter. Capsule size alternation was analyzed by ANOVA on set of >120 cells. (* p < 0.05, ** p < 0.05, ∗∗∗p < 0.001). The red dotted line indicates the average, relative capsule size for the WT strain.

Incubation of encapsulated cryptococcal cells with the cell wall staining (UVtex-2b) and reducing end probe (HA-488) allows for visual localization of the GXM-reducing ends fungal cell wall [63]. Among the tested mutant strains there was no visible change in the reducing end probe signal pattern when compared to the WT strain, consistent with the notion that none of the proteins of interest has modifying the position of polysaccharide-reducing ends (Fig 6A). Immunostaining analysis of the capsule structure of the released polysaccharide-related proteins mutants with mAb 18B7 to GXM presented no visible changes in the peripheral structure of the capsule among most tested strains (Fig 6B). Detailed visual inspection revealed that strains lacking *exa2* (CNAG_05893) and *exa4* (CNAG_07638) presented minor abnormalities of the surface shape of the capsule structure in 8.8% and 10.8% of the tested cells respectively, providing additional evidence of compromise capsular structure in those two mutant strains.

**Fig 6.**
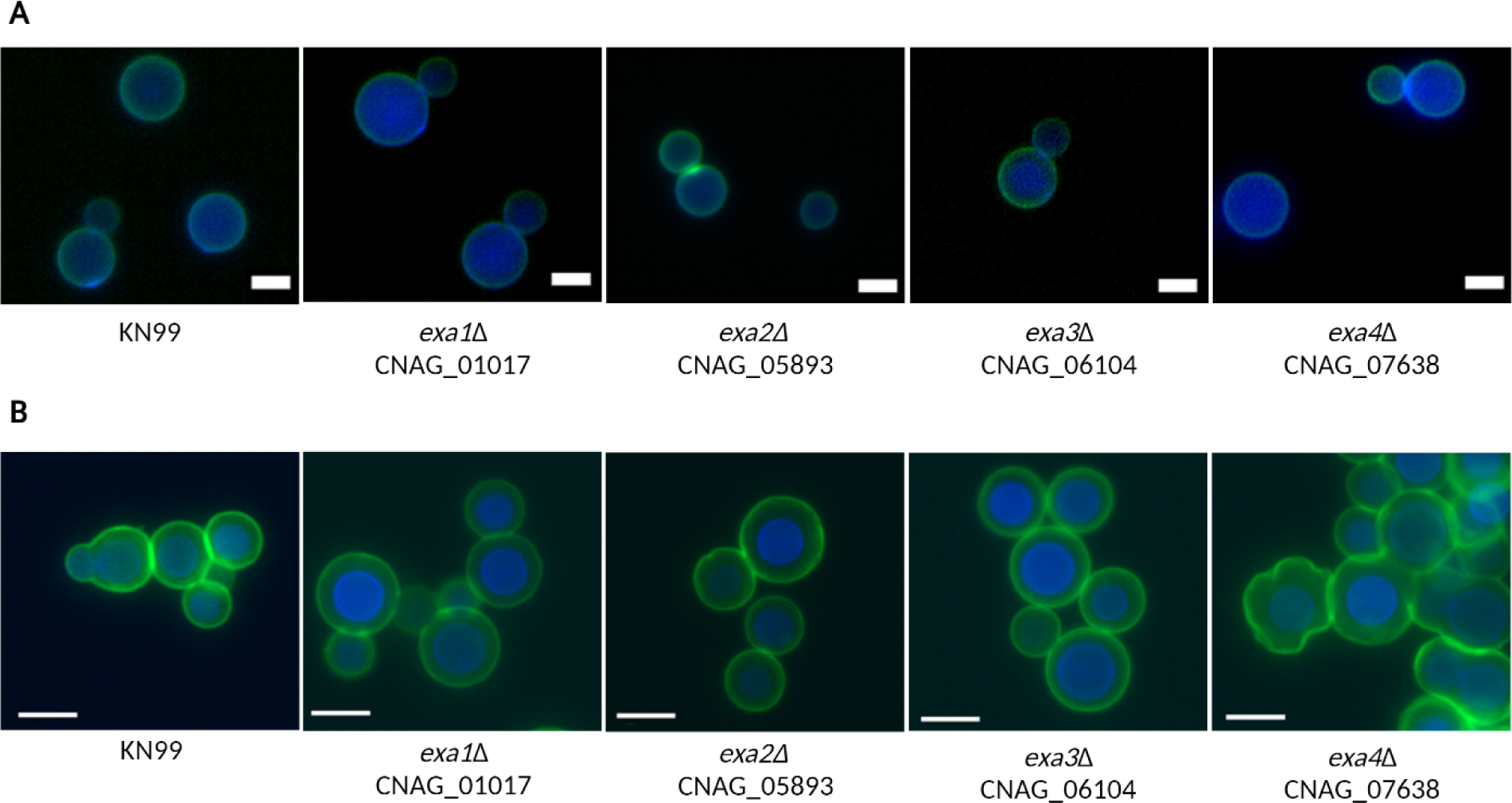
(**A**) Localization of reducing end of capsular polysaccharides. Staining of the capsular polysaccharides reducing ends with R. E. probe (green) confirmed its undisturbed localization within proximity of the cryptococcal cell wall (UVtex 2B) in WT and all tested deletion mutant strains. (**B**) Immunofluorescence imaging of cryptococcal capsule with mAb 18B7 (green) with the cell wall is dye UviTex2B (blue). Green halo corresponds to the edge of the capsule. Strains lacking Spp2 (CNAG_07638) and Spp5 (CNAG_05893) presented minor abnormalities of the surface capsule structure in 10,8% and 8,8% of cells respectively (n>130 cells). Scale bar, 5 µm.

### Alternation of EPS size and secretion efficiency in the SPP deletion mutant strains

To investigate whether the EPS-related proteins had a role in EPS size distribution, we examined the size of EPS polymers released from the WT strain and each of the protein-deficient strains using dynamic light scattering (DLS). Strain *exa3*Δ released EPS with a similar size to those released by the WT strain, with an average size between 108-125 nm. The *exa2*Δ strain released smaller EPS particles (44 nm) than the WT strain. In contrast, EPS particles released by the strain *exa1*Δ were larger, with an average size of 147 nm. The strain lacking *exa4* released two populations of EPS particles 38 nm and 157 nm (Fig 7). Next, we examined the impact of single gene deletion of EPS-related proteins on the efficiency of EPS release. The ELISA assay revealed that the absence of any of the 4 EPS-related proteins (Exa1, Exa2, Exa3, and Exa4) resulted in a significant increase in EPS production (Fig 8).

**Fig 7.**
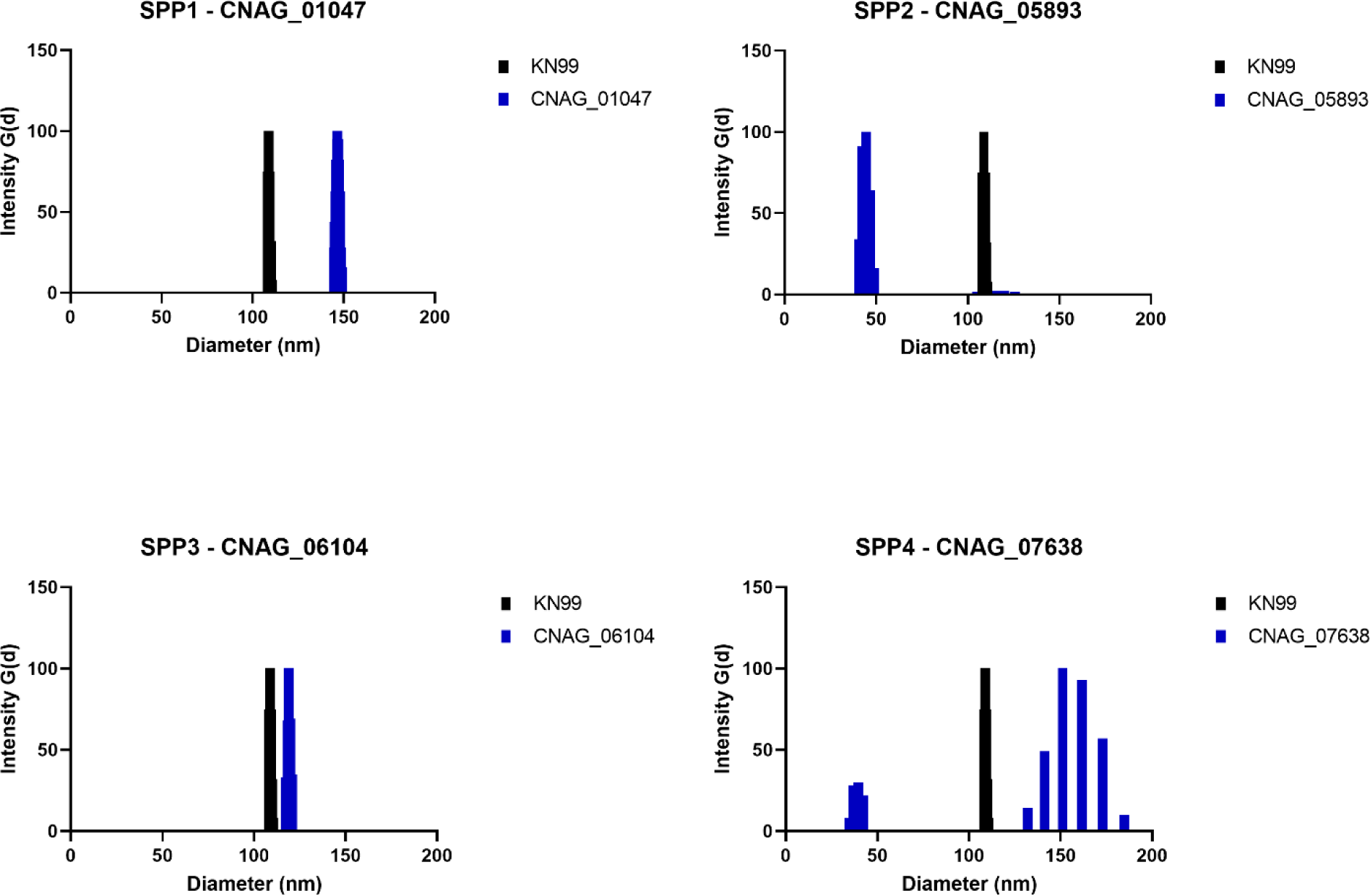
Multimodal size distribution analysis of cryptococcal extracellular polysaccharide fractions harvested from the wild type (KN99) and selected mutant strains.

**Fig 8.**
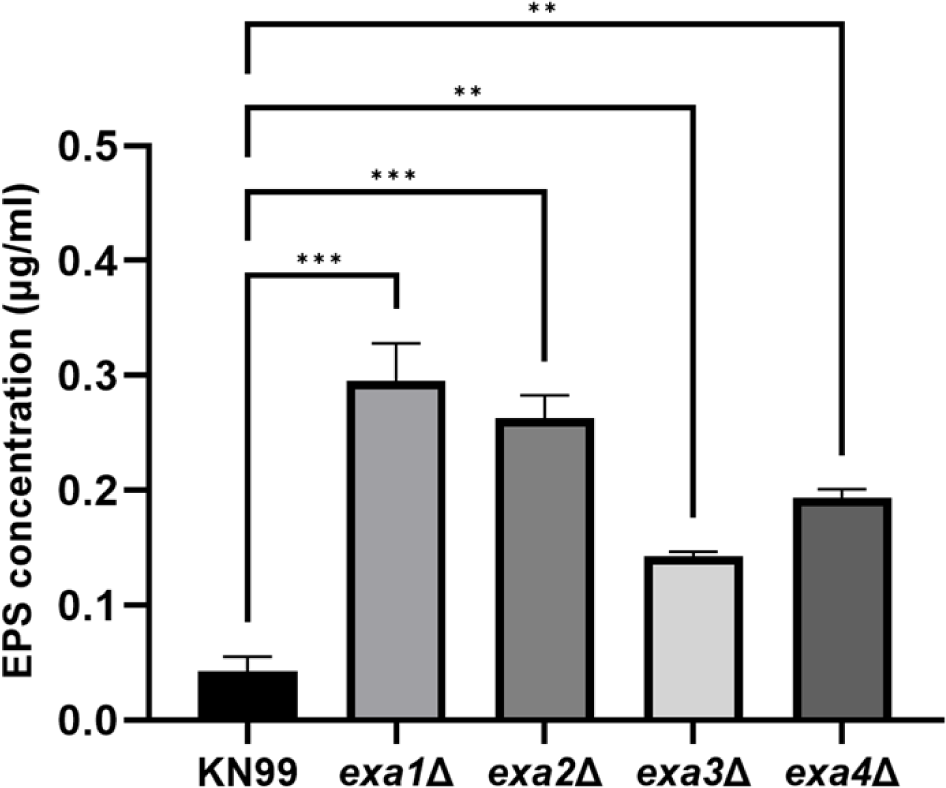
Measurement of the EPS secretion in PBS solution for *C. neoformans* WT (KN99) and selected mutant strains. Mean values were compared with KN99 strain results using an ANOVA test. (* p < 0.05, ** p < 0.05, ∗∗∗p < 0.001).

### The role of EPS-related proteins in biofilm formation

Cryptococcal biofilms are complex structures composed of yeast cells surrounded by polysaccharide matrix and extracellular proteins [24,30,64]. To analyze the function of selected polysaccharide-related proteins we utilized colorimetric XTT and Crystal violet staining assays to observe the metabolic activity and matrix, respectively, of cryptococcal deletion mutant cells within the biofilm. The biofilms of the *exa1*Δ and *exa3*Δ strains showed significant increases in biofilm metabolic activity compared to the WT and other tested strains (Fig 9A). Crystal violet assay confirmed the significant increase in biofilm production in *exa1*Δ and *exa3Δ* strains (Fig 9B). Additionally, the strain lacking *exa2* gene (CNAG_05893) also presented a significant increase in the biofilm matrix formation.

**Fig 9.**
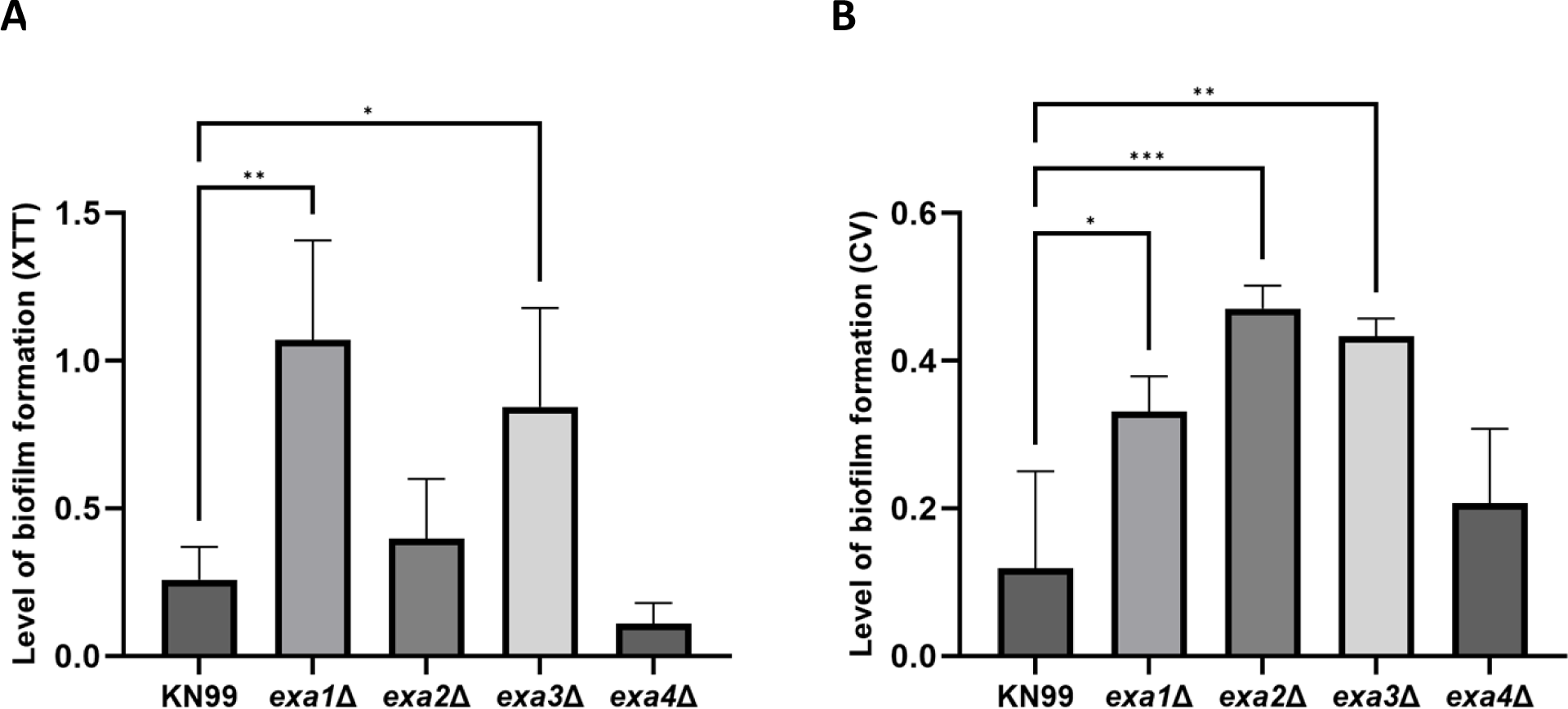
Analysis of the biofilm formation in the tested strains of *C. neoformans*. (**A**) Measurement of the metabolic activity of *C. neoformans* biofilms using an XTT reduction assay. (**B**) Analysis of the cryptococcal biofilm using crystal violet staining assay. The Y axis represents the absorbance values measured at 492 nm for XTT assay and 595 nm for CV assay. Mean values were compared with KN99 strain results using an ANOVA test (* p < 0.05, ** p < 0.05, ∗∗∗p < 0.001).

### Macrophage phagocytic index and virulence in *Galleria mellonella* model

Selected deletion mutant strains were analyzed for phagocytosis by J774.16 macrophage cells and virulence in *G. mellonella*. Elimination of the selected EPS-related (Exa1-Exa4) proteins from cryptococcal cells did not have a significant impact on the efficiency of macrophage phagocytosis measured by analysis of macrophage phagocytic index (data not shown). Reduction of cryptococcal capsule size is often associated with lessened fungal resistance to the innate immune system present in the host [65,66]. We tested the virulence of several SPP deletion mutant strains on invertebrate infection model host *Galleria mellonella* [67]. Inoculation of the *Galleria* larvae with KN99 (WT) strain resulted in the death of all animals within 9 days. Among the tested strains, larvae inoculated with the *exa3*Δ mutant strain died within a similar time frame to WT-infected larvae indicating no major change in their virulence (Fig 10). Most larvae infected with *exa1*Δ and *exa4*Δ survived similar times (P-value 0.038 and P-value 0.049 respectively) to WT-infected hosts, implying only a limited decrease of the cryptococcal virulence. The remaining deletion mutant strain *exa2Δ* presented a substantial reduction of virulence in the *Galleria* model suggesting its important role in fungal pathogenesis. All the phenotypic changes observed with the exa mutant strains were summarized in Table 4.

**Fig 10.**
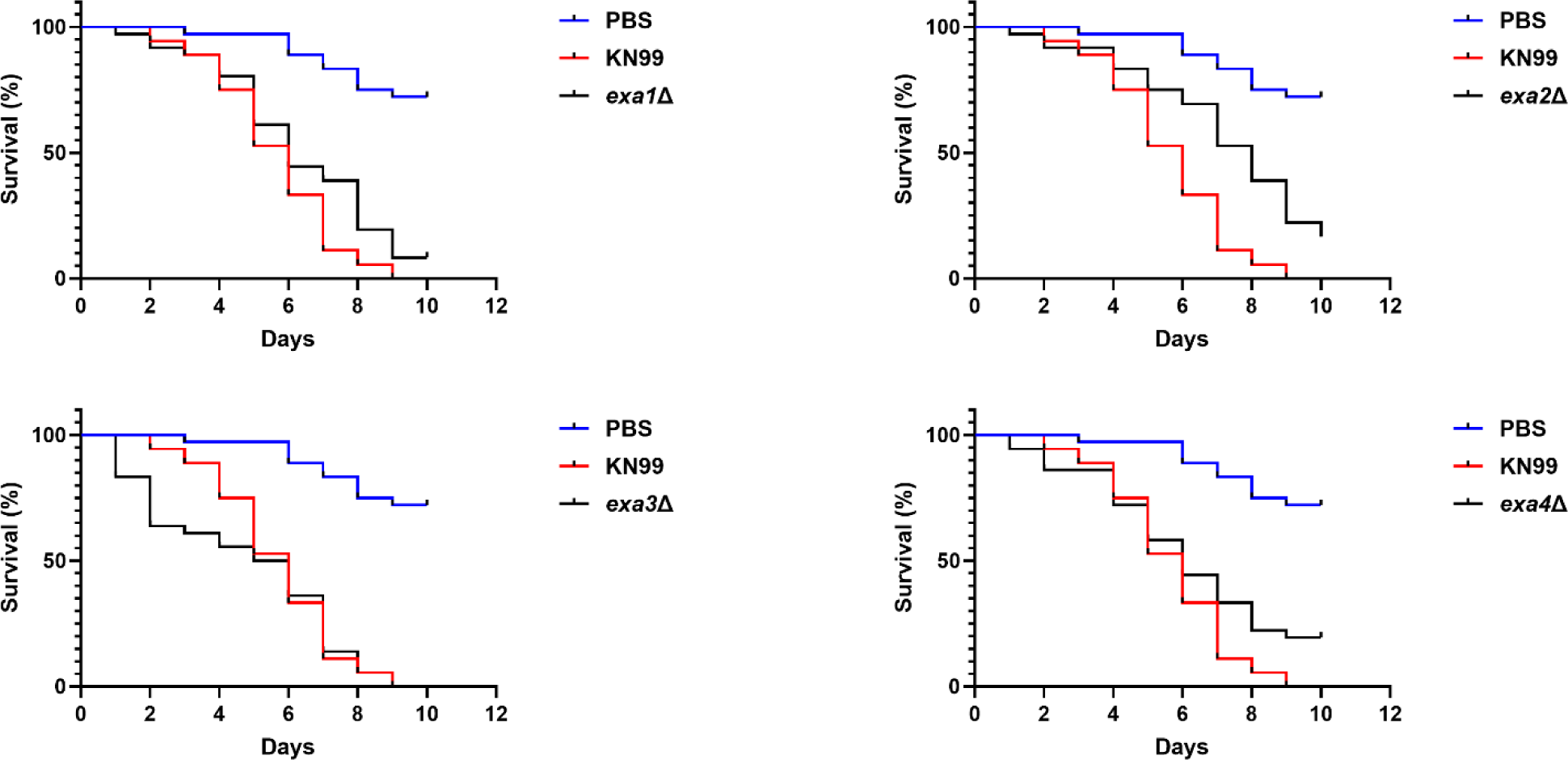
Analysis of the virulence of the selected gene deletion mutants in the invertebrate host infection model *Galleria mellonella*. All tested deletion strains were tested simultaneously and compared to the negative (PBS) and positive (KN99) control, which explains why these curves are the same for each panel. Each graph represents survival of *G. mellonella* infected with each cryptococcal deletion strains and the controls.

**Table 4.**
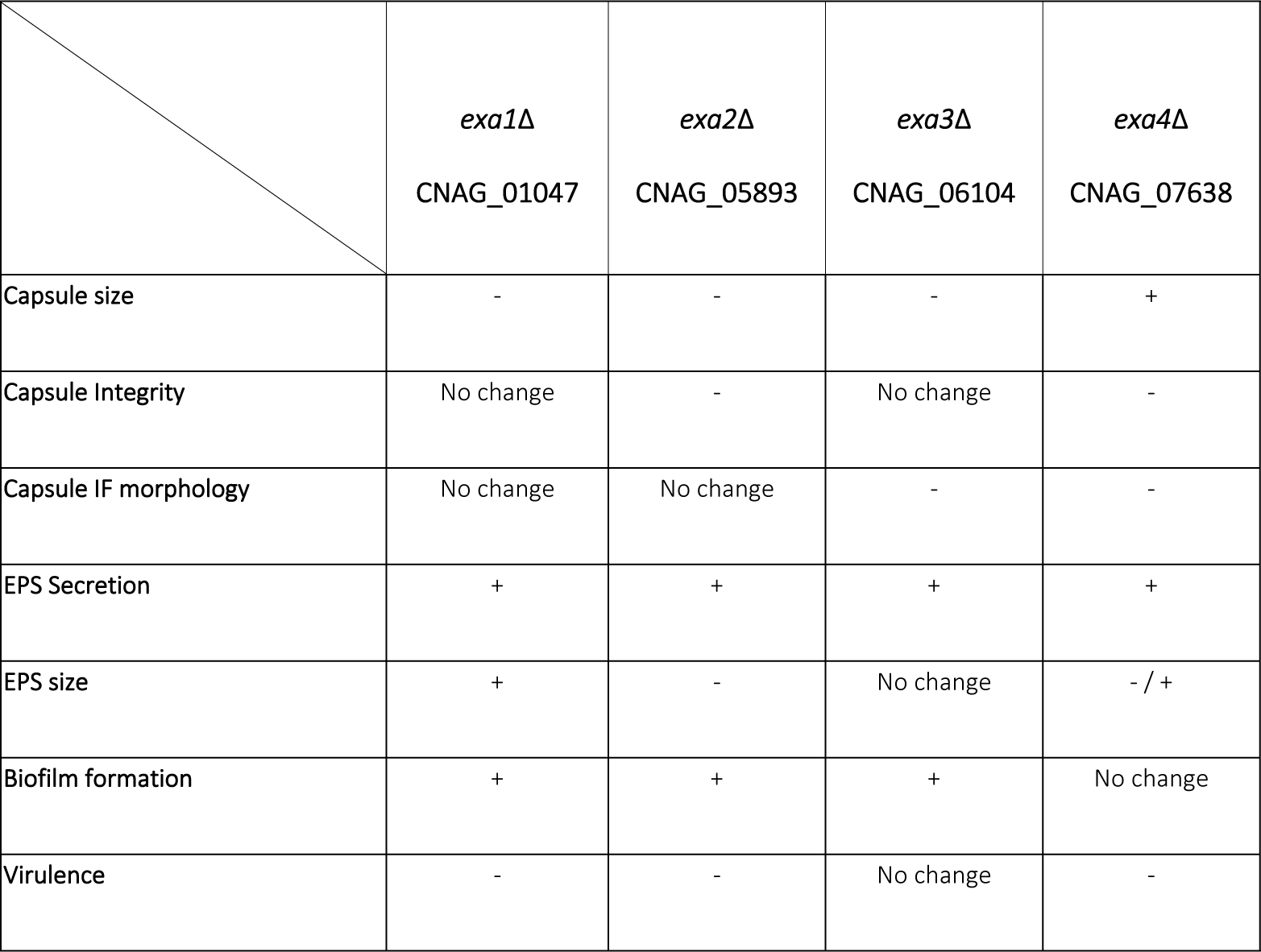
A summary of the findings observed in selected and tested strains.

## DISCUSSION

In this study, we analyzed EPS for associated proteins for the two most clinically relevant species of the *Cryptococcus* genus after placing cells in conditions that stimulated the growth of polysaccharide capsule and secretion of cryptococcal EPS [68,69]. Our mass spectrometry-based analysis identified 62 polysaccharide-associated proteins across 5 cryptococcal strains. For each strain PAGE and proteomic analysis detected a different composition of EPS-associated proteins. Of these 62 proteins, only 22 were identified in both cryptococcal species. A substantial fraction (56%) of the proteins found in our study were previously identified in the proteomic studies of cryptococcal EVs suggesting that these may come from EVs in the preparation, a finding that is also consistent with the proposed role of EVs in the transport of EPS and EPS-related proteins. Sequence comparative analysis of these proteins suggested that many are involved in the carbohydrate metabolic and oxidation-reduction processes. In our EPS-associated protein set, 22 had at least one carbohydrate-acting protein domain. The most common protein domain was the Carbohydrate-Binding Module Family 13 (CBM13)[70,71]. CBM13 domain was first characterized in plant lectin proteins, with affinity to bind sugars including those that compose cryptococcal polysaccharide capsule, such as xylan and galactose [49,72]. Among proteins identified in cryptococcal EPS at least 12 proteins had functions associated with cell wall maintenance. Several of those proteins including chitin deacetylase Cda1 and Cda3, glucan beta glycosidase Ebg1, Eg104, and glucan alpha glycosidase Agn1 have been previously characterized as important proteins for cryptococcal cell wall integrity [33,34,73,74]. Additionally, 16 other proteins had predicted GPI-anchor in their sequence, suggesting a possible connection between cryptococcal capsular and extracellular polysaccharides with cell wall structure [75].

The assembly of the cryptococcal polysaccharide capsule is a complex and not well-understood process [10,60]. The polysaccharide capsule is a major virulence factor and an important barrier protecting cryptococcal cells from external stress factors [6,7,10]. Similarly, to EPS, the capsule is composed of GXM and other non-polysaccharide components that include lipids and proteins [6,7,39]. In our study, we followed up the identification of EPS-associated protein sets with investigations about their possible roles using gene-deletions mutants. A group of 19 proteins with unknown or unspecified functions found in our analysis were selected for evaluation of their function in capsule assembly by phenotypically characterizing protein-deficient mutants from a deletion library (Table 2). Among 11 selected deletion strains, 4 manifested alterations in the capsule size. One strain named *exa4*Δ showed a significant increase in the capsule size, while the other three strains *exa1*Δ, *exa2*Δ, and *exa3*Δ presented cells producing significantly smaller capsule. Further analysis of the selected strains associated the production of Exa2 and Exa4 with the maintenance of the size and structural integrity of the capsule (Fig 5). Additionally, Exa2, Exa3, and Exa4 are predicted to have GPI-anchor in their structure which indicates their localization at the cell wall–capsule region. Interestingly, the absence of *exa2* had a negative impact on capsule size, but significantly increased the integrity of the structure suggesting its potential role in the later stage of capsule remodeling. Furthermore, Exa2 was previously reported in the proteomic studies of cryptococcal EVs [44]. Finally, the visual analysis of the polysaccharide-reducing end patterns showed no difference in fluorescence pattern and was thus not informative regarding a possible role for the tested proteins in this process. Strains lacking *exa3* released EPS of similar size to those released by WT, while EPS molecules released by strains *exa1*Δ were larger. The *exa4*Δ strain released EPS with two distinct sizes. The EPS polymers from *exa2*Δ were approximately 2.5x smaller than those from the WT strain. This atypical EPS release led us to investigate the role of these genes in the regulation of biofilm formation using the XTT reduction assay and crystal violet colorimetric assay [76,77]. The XTT assay indirectly measures the activity of microbial cells integrated into the biofilm matrix by analyzing the reduction of water-soluble formazan in viable fungal cells. Crystal violet stain dyes negatively charged molecules present in the biofilm matrix and on the surface of the cells [78]. Enhanced biofilm formation in *exa1*Δ, *exa2*Δ, and *exa3*Δ strains positively correlated with increased EPS production. The combination of increased EPS release, increased biofilm formation and reduced capsule size suggests a that cryptococcal capsule assembly may be impaired in these strains. The presence of glycosyl hydrolase family 79 domain in Exa3 and the relatively small size of Spp1 (14 kDa) suggests potentially different functions of both proteins in the process of EPS release and capsule formation.

The virulence of selected strains deletion strains was evaluated in *Galleria mellonella,* where most of the tested strains manifested moderate to severe reduction of virulence, as measured by longer survival. The changes in virulence among tested samples could be related to the alterations in capsular size and structure as well as alterations of the EPS secretion. The smallest reduction of virulence was observed in the strains manifesting a significant increase in biofilm production.

In conclusion, our study identified a large set of proteins found in the EPS isolates, and analysis of a subset strongly suggests that some have important roles in the regulation of capsule growth, EPS release, and biofilm formation (Table 3). The presence of many proteins with potential roles in polysaccharide metabolism in the extracellular space supports the notion that the final assembly of these important virulence factors occurs in the extracellular space. We described four novel proteins whose absence was associated with capsular abnormalities suggesting a role in capsule formation. Further analysis of proteins related to capsular and extracellular polysaccharides is essential to better understand the mechanisms of capsular assembly and maintenance. The changes resulting from the loss of the EPS-related proteins impacted major cryptococcal virulence factor followed by a significant decrease in virulence. Our results connect to prior immunological studies showing that a cryptococcal culture filtrate known as CneF elicited strong cell-mediated responses and our findings suggest that some of the proteins described here could have been responsible for those antigenic effects [21,79–81]. Although studies with CneF have not been pursued for more than two decades our results suggest an explanation for how a preparation that contained polysaccharides elicited strong cell-mediated responses. Additionally, the lack of homology to human proteins suggests that those proteins could be considered as a promising potential drug target and/or new antigens for the design of vaccines against *C. neoformans*.

## MATERIALS AND METHODS

### Fungal Growth and Capsule induction

*C. neoformans* strains (H99, Mu-1, 24067) and *C. gattii* strains (409 and KT24066) cells, as well as the single gene deletion mutant cells, were maintained in −80°C freezer stock and grown for study by subculturing in liquid yeast extract peptone dextrose (YPD) media for 2 d at 30° C. Cryptococcal single gene deletion mutant strain libraries were generated by Hiten Madhani at UCSF and purchased from the Fungal Genetic Stock Center (FGSC) [82].

Capsule growth was induced in minimal media (7.5 mM glucose, 10 mM MgSO4, 29.4 mM KH2PO4, 6.5 mM glycine, and 3 µM thiamine-HCl, pH 5.5) for 3 days at 30°C, on the rotor (180 rpm).

### Validation of gene deletion

PCR validation was used to demonstrate NAT Cassette insertion (∼1.5 kb) at each locus of selected deletion mutant strains. Genomic DNA was prepared using CTAB DNA extraction method. PCR reactions were performed using Phusion polymerase and run on a 1% agarose TAE gel at 80v with a 1 kb + ladder. PCR validation was used to demonstrate NAT Cassette insertion (∼1.5 kb) at each locus of the tested deletion mutant strains (S Fig 1).

### Protein and Exopolysaccharide Isolation

The supernatant was isolated from cells by centrifugation (4,000 x g, 15 min, 4°C) and later sterilized using a 0.45 µm filter. Filtered supernatant samples were fractionated with 100 kDa and 10 kDa MWCO Amicon filters for 20 min at 4000 x g. The fraction utilized is the flow through of the 100 kDa filter that was retained by the 10 kDa filter, therefore 10-100 kDa. The retained fractions were stored at 4°C and used in further experiments.

### SDS PAGE gel analysis

For protein gel analysis, 5 µl of the EPS sample was mixed with 2.5 µl of NuPAGE® 4X LDS Sample Buffer, 1 µl of NuPAGE® 10X Reducing Agent, and 1.5 µl of autoclaved MilliQ water. Proteins in the samples were denatured by 10 min incubation at 80°C. For sample analysis 8µl of each sample were loaded on to a 4-20% Mini-PROTEAN® TGX™ Precast Protein Gel and placed in the Mini-PROTEAN Tetra Vertical Electrophoresis chamber. Protein gel electrophoresis was conducted for approximately 2.5h at 70V. Visualization of the protein bands was performed with Pierce Silver Stain Kit (Thermo Scientific) according to the manufacturer’s instructions.

### Mass Spectrometry Sample Preparation and Analysis

To prepare set of proteomic samples for Mass Spectrometry analysis proteins were first reduced with 10 uL of 50 mM dithiothreitol for 1 h at 60°C. Samples were then cooled to RT and pH of samples was adjusted to ∼7.5, followed by subsequent alkylation with 10 uL of 100 mM iodoacetamide in the dark at RT for 15 min. Proteins were digested in solution with Trypsin/LysC (Pierce) at 37°C overnight. Peptides were desalted on a C-18 Oasis Plate and dried in a speedvac concentrator. Peptides were analyzed by reverse-phase chromatography-tandem mass spectrometry on an EasyLC nano HPLC interfaced with a Q-Exactive+ mass spectrometer (Thermo Fisher Scientific). Peptides were separated using a 0%–100% acetonitrile in 0.1% formic acid gradient over 120 min at 300 nl/min. The 75 µm x 25 cm column (ESI Source Solutions) was packed in house with ReproSIL-Pur-120-C18-AQ (3 µm, 120 Å bulk phase, Dr. Maisch). Survey scans of precursor ions were acquired from 350-1800 m/z at 70,000 resolutions at 200 m/z, automatic gain control (AGC) of 3 x 10^6^, and an RF lens setting of 65%. Precursor ions were then individually isolated within 1.6 m/z by data-dependent acquisition with a 15s dynamic exclusion, the top 15 precursors were selected and later fragmented using an HCD activation with a collision energy of 28. Fragment ions were analyzed at 35,000 resolution AGC of 5 x 10^3^ and an intensity threshold of 3.3 x 10^4^. All data files were analyzed using Mascot (Matrix Science, London, UK; version 2.8.2) with the RefSeq_Cryptococcus_neoformans_221213 database (21,284 entries) assuming the digestion enzyme trypsin. Mascot was searched with a fragment ion mass tolerance of 0.02Da and a parent ion tolerance of 5.0 PPM. Peptide identifications from the Mascot searches were processed and imported into Scaffold (Proteome Software Inc.), with peptide validation (5% false-discovery rate) and protein inference (99% confidence) by PeptideProphet [83,84]. Identification of correct cryptococcal protein homologs was performed using BLASTp tool (https://blast.ncbi.nlm.nih.gov/Blast.cgi) in reference to four reference strains (*C. neoformans* var. grubii H99, *C. neoformans* var. *neoformans* JEC21, *C. gattii* VGII R265, and *C. gattii* VGIII CA1280) (Supplementary table 1). The table includes raw MS data and assigned, organized corresponding proteins with the level of protein percent identity of identified homologous proteins among selected reference cryptococcal strains. In Supplementary Table 1 we assigned a confidence level to each of the identified proteins in the organization tab. Proteins that were identified with high confidence (based on two unique peptides from at least 2 out of 3 biological replicates for the same strain with at least two peptides in each replicate) were marked with green color while proteins identified with medium confidence (based on unique peptides from at least 2 out of 3 biological replicates for the same strain with at least one peptide in each replicate) were marked with yellow color. Only proteins present in at least two biological replicates with at least one unique peptide were selected for further analysis. Proteins were further categorized into groups based on their presence in both species *C. neoformans* and *C. gattii*, only in *C. neoformans* or only in *C. gattii*. In the segregation tab proteins were categorized into groups based on their presence detected in EPS samples in both *C. neoformans* and *C. gattii* complex (blue), only in *C. neoformans* complex (yellow), or only in *C. gattii* complex (grey).

### Capsule size measurement

The cryptococcal capsule of the WT and selected deletion mutant strains was performed using the India ink negative staining method. Images were taken on an Olympus AX70 microscope were then analyzed using the Quantitative Capsule Analysis program or manually using ImageJ software [85]. The relative capsule size was established as a cell capsule-to-cell body diameter ratio. Significance was determined using an ANOVA on GraphPad Prism software.

### Microscopic observation of capsule morphology

To observe the capsule cryptococcal cells were washed two times in the blocking buffer (1% BSA) and cell density was adjusted to 10^7^ cells/ ml. Cells were incubated with 50 μg/ml of mAbs 18B7 for 1 h at room temperature and after incubation washed twice with 500 μl of PBS and, further incubated for 1.5h with goat anti-mouse IgG1 FITC-conjugated antibody (5 μg/ml). In the last 10 min of incubation, Uvitex-2b was added to cell suspension to visualize the cryptococcal cell wall. After incubation cells were washed two times with PBS. Cells were analyzed using the Olympus AX70 microscope and ImageJ software.

To analyze the localization of the capsular polysaccharide-reducing ends cryptococcal encapsulated cultures were washed two times in the blocking buffer (1% BSA) and adjusted to 10^7^ cells/ ml. The cells were then incubated overnight at room temperature with the 3 μm fluorescent reducing-end probe [63]. During the last 10 minutes of incubation, Uvitex-2b was added to cell suspension to visualize the cryptococcal cell wall. After incubation cells were washed two times with PBS and analyzed using the Olympus AX70 microscope and ImageJ software.

### *In silico* pProtein domain and homology analysis

Sequences of the proteins identified in this study were analyzed with the InterPro software (https://www.ebi.ac.uk/interpro/) and CAZY database to identify the potential function of protein domains. The protein interaction network was organized and arranged using STRING protein network (VER 11.5) software. The network was generated using default settings, with a selection of medium confidence. Sequences of *C. neoformans* proteins, found in this study, were obtained for the Fungal database (Fungi DB) and analyzed with the BLASTP software (blast.ncbi.nlm.nih.gov) to find homologous proteins in the proteome of Human and major fungal pathogens including *C. albicans*, *H capsulatum* and *A. fumigatus*[86]. A similarity between the analyzed proteins was indicated as a percentage value and presented in the form of a table.

### EPS analysis

Encapsulated *C. neoformans* cells were washed 3 times with sterile PBS and 10^6^ cells were transferred to 1 ml of PBS and left in the 30°C incubator for 24 h. After the incubation samples were centrifuged to pellet the cells and 1 ml of the supernatant was used in the ELISA assay to determine the concentration of the EPS. EPS concentration was established using anti-GXM IgG 18B7 at 5 µg/ml 50 µl/well and goat anti-mouse IgG1 + alkaline phosphatase (Southern Biotech) in the 96-well plate (CORNING). The assay was read at 405 nm with EMAX® Plus Microplate Reader and SoftMax Pro Software. Significance was determined using an ANOVA on GraphPad Prism software.

Measurement of the EPS particle size distribution was performed with a Zeta Potential Analyzer (Brookhaven Instruments). A 100 µl of the EPS sample was placed in the Cuvette (Eppendorf) at room temperature and placed into the machine. The measurements were taken for the 10 runs repeated of 1m each. The multimodal size distributions were presented as a graph to visualize differences among the samples.

### Capsule integrity assay

Encapsulated WT and selected mutant strains were washed with PBS and brought to the density of 5×10^7^ cells/ml. The capsule integrity challenge was performed on ice using a horn sonicator machine (Fisher Scientific Sonic Dismembrator F550 W/ultrasonic Convertor) on power settings 7 (17W) for 15 sec to disturb the outer region of the capsule [87]. The capsule size of each sample was measured via India ink negative staining on an Olympus Light microscope analyzed using the ImageJ software Significance was determined using an ANOVA on GraphPad Prism software.

### Analysis of biofilm formation

To measure biofilm formation 200 µl of 1 x 10^7^ cell/ml of MM cultures of *C. neoformans* were distributed onto two separate polystyrene 96-well plates and incubated at 37°C for 48 h. Cultures were carefully washed three times with 200 µl of PBS and drained in an inverted position to remove residual PBS. The biofilm formation was analyzed using XTT assay and crystal violet staining assay. Metabolic activity of cryptococcal biofilm formation in the tested *C. neoformans* strains was performed with (XTT) 2,3-bis (2-methoxy-4-nitro-5-sulfophenyl)-5-[(phenylamino) carbonyl]-2H-tetrazolium-hydroxide reduction assay. A 96-well plate was filled up with 200 µl per well of XTT salt solution (1mg/ml in PBS) and menadione solution (1 mM in acetone; Sigma) and incubated for 3 h at 37°C. The colorimetric change was measured and analyzed using an EMAX® Plus Microplate Reader and SoftMax Pro Software. Biofilm matrix formation was measured with a crystal violet staining assay. Each of the washed and dried wells was stained with 110 μl of 0.4% aqueous crystal violet solution for 45 min. Next, each well was washed three times with 200 μl of sterile dd water and de-stained with 200 μl of 95% ethanol. After 45 min of de-staining, 100 μl of de-staining solution was transferred to a new 96-well plate and measured with a microtiter plate reader EMAX® Plus Microplate Reader and a SoftMax Pro Software at 595 nm. Significance was determined using an ANOVA on GraphPad Prism software.

### Macrophage phagocytic index

The phagocytic index was performed based on a published protocol [88]. Briefly, J774.A1 cells (ATCC, Gaithersburg, MD-US) were seeded in complete media, composed by DMEM (Cytiva, Marlborough, MA-US) and 10% FBS (R&D Systems) in CO_2_ incubators at 37 °C and 5% CO_2_. The cryptococcal cells counted for a final concentration of 1.5 x 10^6^ per well in activated media and incubated with 20% of guinea pig serum complement for 1 h at 30°C. Next, cryptococcal cells were transferred to the J774.A1 cells and plates were incubated for 2 h. After incubation cells were washed twice with PBS and the phagocytic index was determined by the number of internalized Cryptococcal cells divided by the total number of macrophages. Significance was determined using an ANOVA.

### Galleria mellonella virulence assay

The assay was performed as described in this publication [67]. *Galleria mellonella* larvae were selected and used for the following experimental groups (12 larvae per group) and infected with selected strains of *C. neoformans*. Each larva was injected with 10μl of the inoculum (10^4^ cells/µl) into the last right proto leg using a U-100 insulin syringe (27G). After inoculation, larvae were kept in separate Petri dishes and incubated at 30°C. The viability of the larvae was assessed daily based on change of color and response to touch. Significance was determined using a Log-rank Mantel-Cox test on GraphPad Prism software.

## ACKNOWLEDGMENTS

Special thanks go to Robert N. Cole and Lauren DeVine in the Spectrometry and Proteomics Facility at Johns Hopkins School of Medicine for mass spec analysis and data consultation.

